# *pkgndep*: a tool for analyzing dependency heaviness of R packages

**DOI:** 10.1101/2022.02.11.480073

**Authors:** Zuguang Gu, Daniel Hübschmann

**Affiliations:** Molecular Precision Oncology Program, National Center for Tumor Diseases (NCT), Im Neuenheimer Feld 280, 69120 Heidelberg, Germany; Heidelberg Institute of Stem Cell Technology and Experimental Medicine (HI-STEM), Im Neuenheimer Feld 280, 69120 Heidelberg, Germany; German Cancer Consortium (DKTK), Im Neuenheimer Feld 280, 69120 Heidelberg, Germany; Department of Pediatric Immunology, Hematology and Oncology, University Hospital Heidelberg, 69120 Heidelberg, Germany

## Abstract

**Summary:** Numerous R packages have been developed for bioinformatics analysis in the last decade and dependencies among packages have become critical issues to consider. In this work, we proposed a new metric named *dependency heaviness* that measures the number of unique dependencies that a parent brings to a package, and we proposed possible solutions for reducing the complexity of dependencies by optimizing the use of heavy parents. We implemented the metric in a new R package *pkgndep* which provides an intuitive way for dependency heaviness analysis. Based on *pkgndep*, we additionally performed a global analysis of dependency heaviness on CRAN and Bioconductor ecosystems and we revealed top packages that have significant contributions of high dependency heaviness to their child packages.

**Availability and implementation:** The package *pkgndep* and documentations are freely available from the Comprehensive R Archive Network (CRAN) https://cran.r-project.org/package=pkgndep. The dependency heaviness analysis for all 21,741 CRAN and Bioconductor packages retrieved on 2021-10-28 are available at https://pkgndep.github.io/.

## 1 Introduction

Reliable and robust software is essential for data analysis in bioinformatics. In the last decade, R has rapidly become a major programming language for developing software for biological data analysis, including data processing and visualization, statistical modeling, interactive web application and reproducible report generation. It is applied in broad research fields of genomics, transcriptomics, epigenomics, proteomics, metabolomics and more. The reusable and extensible code are normally organized in forms of R packages, which are distributed on public repositories such as the Comprehensive R Archive Network (CRAN) and Bioconductor. Publications describing R packages have also been increasing. By 2022-01-28, using search term “(R package [Title]) OR Bioconductor [Title/Abstract]” on PubMed resulted in 2,496 publications. Dependencies always exist in packages where a package imports functionalities from other packages. As the number of R packages increases, dependencies among packages become even more complicated. By 2021-10-28, there are 154,477 direct dependency relations among total 18,325 and 3,416 packages from CRAN and Bioconductor respectively.

In the ecosystems of CRAN and Bioconductor, package dependencies are represented as a large directed graph. Research on the dependency graph helps to reveal interesting patterns of package relations, such as top packages that play leading roles or package modules for specific analysis topics (Mora-Cantallops *et al.*, 2020). There are also R packages for dependency graph visualization, such as *deepdep, pkgnet* and *pkggraph*. They support visualizing subgraphs induced from a single package.

Denote a package as *P*, in the dependency graph, the total packages located in the upstream of *P* compose its dependencies. If *P* depends on a large number of upstream packages, there will be the following risks: 1. Users have to install a lot of additional packages when installing *P*, which would bring the risk that installation failure of any upstream package stops the installation of *P*. 2. The number of packages loaded into the R session after loading *P* will be huge, which increases the difficulty to reproduce a completely same working environment on another computer. 3. Dependencies of *P* will spread to all its child packages if there is any. 4. On platforms for continuous integration such as GitHub Action or Travis CI, automatic validation of *P* could easily fail due to the failures of its upstream packages. *P* inherits all its dependencies from its parent packages. Thus, identifying the parents that contribute high dependencies on *P*, *i.e., P*’s heavy parents, helps to reducing the complexity of *P*’s dependencies.

In this work, we proposed a new metric named *dependency heaviness* that measures the number of additional dependency packages that a parent brings to its child package and are unique to the dependency packages imported by all other parents. We implemented the heaviness metric in an R package named *pkgndep*. Additionally, *pkgndep* provides an intuitive way for visualizing package dependencies. Based on *pkgndep*, we also performed a preliminary analysis of dependency heaviness on CRAN and Bioconductor ecosystems and we revealed top packages that contribute significantly high heaviness on their child packages.

## 2 Methods

Every R package has a DESCRIPTION file for its metadata. In it, its dependency packages are listed in the fields of “Depends”, “Imports”, “LinkingTo”, “Suggests” and “Enhances”. Denote a package as *P*, packages listed in “Depends”, “Imports” and “LinkingTo” are mandatory to be installed when installing *P*. Functions, S4 methods or S4 classes defined in packages in “Depends” and “Imports” are imported to the namespace of *P* according to the rules defined in *P*’s NAMESPACE file. Packages listed in “Suggests” and “Enhances” are not mandatory to be installed. They are optionally used in *P*, such as in vignettes, in examples, or in functions that are only required when the functions are called. Readers please refer to the official R manual “Writing R Extensions” for more details.

R package dependencies can be modelled as a directed graph. For the ease of discussion, the following dependency categories for *P* are defined. 1) **Strong parent packages:** the packages listed in “Depends”, “Imports” and “LinkingTo”. For simplicity, in the text we always refer *parent packages* to strong parent packages. 2) **Weak parent packages:** the packages listed in “Suggests” and “Enhances”. 3) **Strong dependency packages:** the total packages by recursively looking for parent packages. Note all strong dependency packages are mandatory to be installed for *P* and a failure of any of them stops installation of *P*. For simplicity, in the text we always refer *dependency packages* to strong dependency packages. 4) **All dependency packages:** the total packages by recursively looking for parent packages, but on the level of *P*, its weak parents are also included. It simulates when full functionality of *P* is required, or in other words, when all its weak parents are changed to strong parents, the number of strong dependency packages it requires. 5) **Child packages:** the packages whose parent packages include *P*. These are the packages on which *P* has a direct contribution of dependencies.

Next the heaviness from a parent is defined. If package *A* is a parent of *P*, the heaviness of *A* on *P* denoted as *h* is calculated as *h* = *n*_1_ - *n*_2_ where *n*_1_ is the number of strong dependencies of *P*, and *n*_2_ is the number of strong dependencies of *P* after changing *A* from a strong parent to a weak parent, *i.e.*, moving *A* to “Suggests” of *P*. In other words, the heaviness measures the number of additionally required strong dependencies that *A* brings to *P*, and these dependencies are not imported by any other parent.

When *A* is a weak parent of *P*, *n*_2_ is defined as the number of strong dependencies of *P* after changing *A* to a strong parent of *P*, *i.e.*, moving *A* to “Imports” of *P*. In this scenario, the heaviness of the weak parent is calculated as *h* = *n*_2_ - *n*_1_.

Note for some (*A*, *P*) pairs, it is possible *n*_1_ = *n*_2_ which results in *h* = 0, *i.e.*, moving parent *A* to “Suggests” of *P* won’t reduce the number of *P*’s strong dependencies. It mostly occurs when a second parent, denoted as *B*, also depends on *A* (directly or indirectly). Thus, only moving *A* to “Suggests” of *P* won’t affect that *A* is still imported as a strong dependency by *B*. For example, package *GenomicRanges* is a parent of package *bsseq* where *GenomicRanges* is a general-purpose package for dealing with genomic intervals. *bsseq* has a second parent package *BSgenome* that also depends on *GenomicRanges*. Thus, moving *GenomicRanges* to “Suggests” of *bsseq* won’t change the number of strong dependencies for *bsseq* because *BSgenome* still imports it. As a result, *GenomicRanges* contribute zero heaviness on *bsseq* as its parent.

## 3 Results

The dependency heaviness analysis is represented as a customized heatmap. Figure 1 demonstrates the dependency heatmap for package *ComplexHeatmap*. On the heatmap, *ComplexHeatmap*’s strong and weak parents are rows split by their dependency relations, *i.e.*, “Depends” or “Imports”. The dependency packages imported by each parent are columns and they are split into two groups of base packages and other packages. The red and blue cells correspond to dependency packages that each parent requires. Total strong dependencies required for *ComplexHeatmap* are marked by purple lines on both rows and columns. On the right side of the dependency heatmap are three additional bar plot annotations. The first bar plot annotation illustrates numbers of functions or classes imported from parents, which are parsed from *ComplexHeatmap’*s NAMESPACE file. The bar plot can distinguish the following different scenarios of imports for each parent: 1. The whole namespace is imported; 2. A limited number of functions are imported; 3. A limited number of S4 methods/classes are imported (*e.g.*, in package *biovizBase*); 4. The whole namespace excluding a limited number of functions is imported (*e.g.*, in package *dplyr*); 5. No function is imported (*e.g.*, in package *ggplot2*). The second bar plot annotation illustrates the number of required packages for each strong or weak parent, which is simply the number of hits on the corresponding row. The third bar plot annotation illustrates the heaviness of each strong or weak parent package on *ComplexHeatmap*, which is the number of required packages uniquely imported by the parent compared to all other strong parents.

**Fig. 1.**
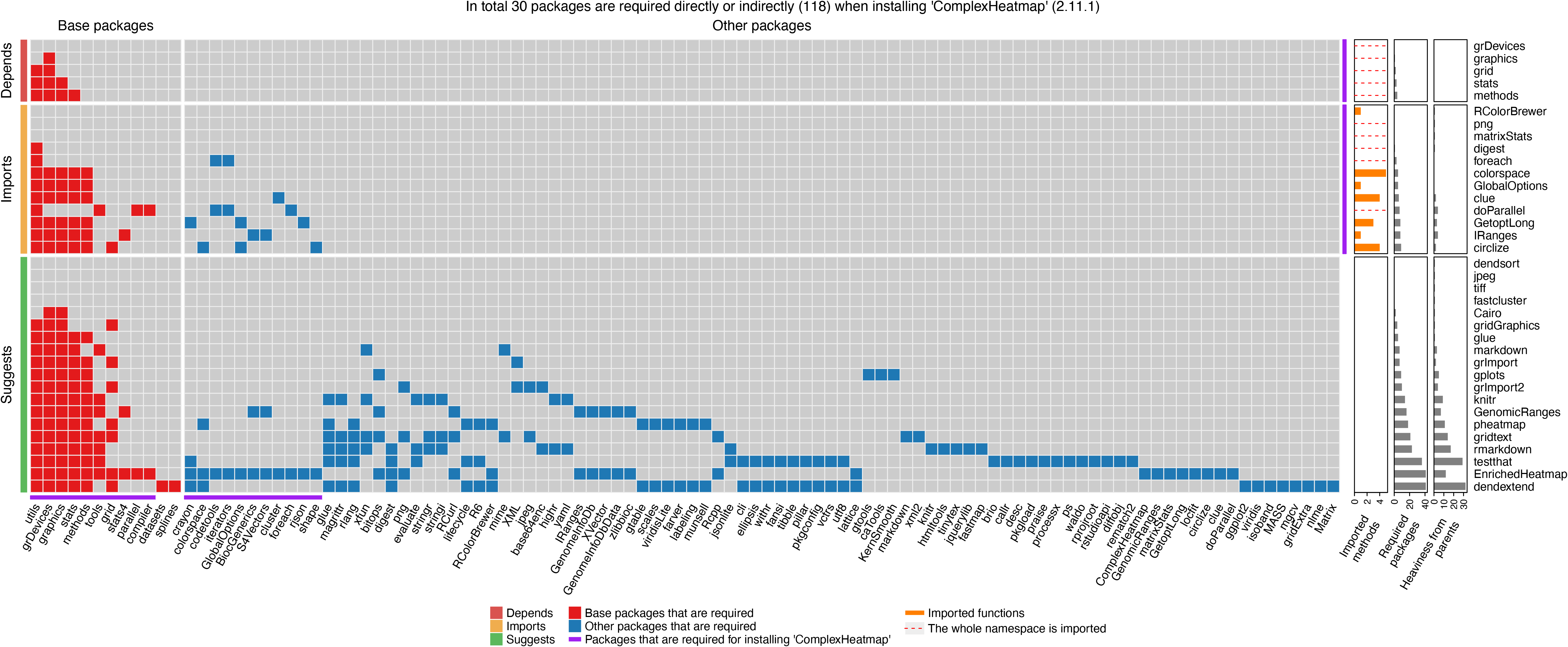
Dependency heaviness analysis on package *ComplexHeatmap*. The interpretation of the plot can be found in the main text.

As illustrated in Figure 1, *ComplexHeatmap* has 30 strong dependency packages, while if including dependencies from all weak parents as well, the number will dramatically increase to 118 (almost 4 times high). Since we are also the developers of *ComplexHeatmap*, during the development we moved some heavy parents, which only provide enhanced functionalities for *ComplexHeatmap* but are not expected to be frequently used, to “Suggest”, such as the package *dendextend* (contributing a heaviness of 32) which is only used for coloring dendrogram branches, and the package *gridtext* (contributing a heaviness of 14) which is only for customizing text formats in heatmaps. These two packages are only required when the corresponding functionalities are used by users.

The dependency heatmap gives hints for reducing dependency complexity of a package. Generally speaking, if a parent has a high heaviness compared to other parents, an optimization might be considered, especially if only a few functions are imported from the parent. In Supplementary File 1 which contains dependency heaviness analysis for package *mapStats*, an extremely heavy parent *Hmisc* can be observed where *Hmisc* has a heaviness of 49 which is almost 60% of the total number of required packages of *mapStats.* If moving *Hmisc* to “Suggests” of *mapStats*, the number of strong dependencies can be reduced from 83 to 34. The first bar plot annotation shows there is only one function imported from *Hmisc.* A deep inspection into the source code of *mapStats* reveals that a function `capitalize()` from *Hmisc* is imported to *mapStats.* `capitalize()` is a simple function that only capitalizes the first letter of a string. The 49 additional dependencies imported from *Hmisc* can be avoided by simply reimplementing a function `capitalize()` by developer’s own.

In package *pkgndep*, The function `pkgndep()` performs dependency heaviness analysis for a package. Later the function `plot()` generates the dependency heatmap and the function `dependency_report()` generates an HTML report of the complete dependency analysis.

We performed heaviness analysis for all 18,325 CRAN packages and 3,416 Bioconductor packages retrieved on 2021-10-28 (Supplementary File 2, , also available at https://pkgndep.github.io/). We systematically analyzed the heaviness of packages on their child packages. For a package denoted as *P*, the mean heaviness over its children was calculated. This metric also measures the expected number of additional dependencies for a package if *P* is added as its new parent. On top of the list of packages ordered by their heaviness on children (Supplementary File 3), we found there are some popular R packages for bioinformatics analysis having high heaviness on their child packages, such as *Seurat* (85, 31 children), *minfi* (58, 38 children), *WGCNA* (50, 32 children) and *Gviz* (42, 37 children) where the first value in the parentheses is the heaviness. Thus, packages depending on these heavy packages should be aware of the risks from high dependencies they bring in.

## 4 Discussion and conclusion

Heaviness analysis provides hints for reducing the complexity of package dependencies, but how to optimize depends on the specific use of parent packages in the corresponding package. As has been demonstrated in the example of *mapStats*, the heavy parent can be avoided by implementing a same function as the imported one, but this scenario is not common. Common cases are when heavy parents are used in analysis less used by users. This is typical for some packages for bioinformatics analysis where they provide core analysis as well as secondary analysis. Parent packages for secondary analysis might be arranged as weak parents if they are heavy and not expected to be frequently used by users (see an example in Supplementary File 4). Nevertheless, there are still scenarios when reduction of heavy parents could not be performed: 1) When a package extends the functionality of a heavy parents, the parent must be its strong parent. 2) A heavy parent provides core functionality to a package. 3) When S4 methods or S4 classes are imported from a parent package.

The heaviness measures the number of additional dependency packages that a parent uniquely imports. However, there are scenarios when multiple parents import similar sets of dependencies, which results in heaviness for individual parents being very small. In Supplementary File 5 of the dependency analysis for package *DESeq2*, its two parent packages *geneplotter* and *genefilter* import 51 and 53 dependencies respectively, among which 50 are the same. Due to the high overlap, the heaviness of *geneplotter* and *genefilter* on *DESeq2* are only 1 and 2. However, if considering the two parents together, *i.e.*, by moving both to “Suggests” of *DESeq2*, 23 dependency packages can be reduced.*pkgndep* also proposes a metric named *co-heaviness* (by the function `co_heaviness()`) which measures the number of reduced dependencies by simultaneously changing two parents to weak parents (Supplementary File 5). Nevertheless, the co-effect of multiple parents can always be easily observed from the dependency heatmap.

Current studies on package dependencies focused on the global effect of a package on all its child packages, *e.g.*, through the degree centrality in the dependency graph analysis. However, a package with more child packages does not necessarily mean it contributes more dependencies. *E.g.*, package *Rcpp* has 2,662 child packages but it only contributes an average heaviness of 0.6 to them. In this study, we proposed a new metric that helps to find the parent contributing heavy dependencies to a child package. We believe *pkgndep* will be a useful tool for R package developers to properly handle the dependency complexity of their packages.

## References

Mora-Cantallops, M. et al. (2020) A complex network analysis of the Comprehensive R Archive Network (CRAN) package ecosystem. Journal of Systems and Software, 170, 110744.

