## Supplementary File 1 for "*pkgndep*: a tool for analyzing dependency heaviness of R packages": suppl_1_mapStats_report.html

Dependency information for package 'mapStats'


### Dependency analysis for package mapStats

#### General information

|  |  |
| --- | --- |
| CRAN link | link |
| Package version | 2.4 |
| Number of strong dependencies | 83 |
| Number of all dependencies | 83 |
| Number of parent packages | 9 |
| Max heaviness from parent packages | 49 |
| Total heaviness from parent packages | 62 |
| Number of parent packages (including `Suggests` and `Enhances`) | 9 |

#### Dependency heatmap

In the following dependency heatmap, rows are the parent packages of mapStats and columns are the dependency packages that each parent package brings in.
On the right side of the heatmap, there are three barplot annotations: 1. number of imported functions/S4 methods/S4 classes from parent packages; 2. number of
dependency packages from each parent package; 3. heaviness of each parent package on mapStats.

Adjust heatmap size:
increase
decrease
reset

xml version='1.0' encoding='UTF-8' ?


Base packages
Other packages
Depends
Imports
stats
methods
graphics
utils
grDevices
grid
splines
tools
lattice
Matrix
survival
Rcpp
foreign
glue
MASS
stringr
magrittr
stringi
minqa
numDeriv
mitools
DBI
sp
Formula
ggplot2
latticeExtra
cluster
rpart
nnet
gtable
gridExtra
data.table
htmlTable
viridis
htmltools
base64enc
digest
isoband
mgcv
rlang
scales
tibble
withr
png
jpeg
RColorBrewer
knitr
checkmate
htmlwidgets
rstudioapi
viridisLite
fastmap
nlme
farver
labeling
lifecycle
munsell
R6
ellipsis
fansi
pillar
pkgconfig
vctrs
evaluate
highr
yaml
xfun
backports
jsonlite
colorspace
cli
crayon
utf8
e1071
class
KernSmooth
proxy
plyr


0
1
2
3

Imported
methods


0
20
40
60

Required
packages


0
20
40

Heaviness from 
parents
lattice
maptools
survey
RColorBrewer
colorspace
sp
classInt
reshape2
Hmisc
In total 83 packages are required directly or indirectly (83) when installing 'mapStats' (2.4)
Depends
Imports


Base packages that are required
Other packages that are required
Packages that are required for installing 'mapStats'


Imported functions

The whole namespace is imported

  
  

#### Dependency table

"Import" information is from the NAMESPACE file of mapStats.

**imports:** number of imported functions/objects; **importMethods**: number of imported S4 methods; **importClasses**: number of imported S4 classes.

**Required packages:** number of strong dependency packages for each of the parent package (or in other words, number of dependency packages the parent package brings in).

**Heaviness from parent on mapStats:** number of required packages that can be reduced if moving parent package to `Suggests` of mapStats.

| Parent package | Field | imports | importMethods | importClasses | Required packages | Heaviness from parent on mapStats |
| --- | --- | --- | --- | --- | --- | --- |
| **survey** | Depends | The whole set of functions/methods/classes from parent package is imported to the namespace of mapStats. | | | 15 | 5 |
| **maptools** | Depends | The whole set of functions/methods/classes from parent package is imported to the namespace of mapStats. | | | 9 | 1 |
| **lattice** | Depends | The whole set of functions/methods/classes from parent package is imported to the namespace of mapStats. | | | 5 | 0 |
| **Hmisc** | Imports | 1 | 0 | 0 | 67 | 49 |
| **classInt** | Imports | 3 | 0 | 0 | 10 | 5 |
| **reshape2** | Imports | 1 | 0 | 0 | 10 | 2 |
| **RColorBrewer** | Imports | 1 | 0 | 0 | 0 | 0 |
| **colorspace** | Imports | 1 | 0 | 0 | 4 | 0 |
| **sp** | Imports | 1 | 0 | 0 | 7 | 0 |

---

Analysis was done with pkgndep.
