## Supplementary File 2 for "*pkgndep*: a tool for analyzing dependency heaviness of R packages": suppl_2_all_CRAN_Bioconductor_packages.html

Dependency heaviness analysis for all CRAN/Bioconductor packages


### Dependency heaviness analysis for all CRAN/Bioconductor packages

Search package

R packages were retrieved from CRAN/Biocoductor on 2021-10-28, which include **18,325** packages from CRAN and **3,416** packages from Bioconductor (bioc version 3.14).

All the reports were generated by the following code:

```
library(pkgndep)
x = pkgndep("pkg")  # "pkg" is the package name
dependency_report(x)
```

The complete table of all R packages can be found here: https://docs.google.com/spreadsheets/d/1R4ICniRDBuaJlPL6YUN0IzAjmnp73tllPnJeBeLUB98/edit?usp=sharing.
