## Supplementary File 3 for "*pkgndep*: a tool for analyzing dependency heaviness of R packages": suppl_3_heaviness_on_children.html

Heaviness on child packages


### Heaviness on child packages

###### Zuguang Gu

#### 2022-02-02

##### Definition

**Strong parent packages:** The packages listed in the `Depends`, `Imports` and `LinkingTo` fields of the `DESCRIPTION` of a package denoted as *P*. They are also called the strong direct dependency packages of *P*. Strong parent packages are enforced to be installed when installing package *P*. To make it easy to discuss, in the text we always refer parent packages to strong parent packages.

**Strong dependency packages:** The total packages by recursively looking for parent packages. They are also called upstream packages. Note strong dependency packages contain parent packages. Strong dependency packages are enforced to be installed when installing package *P*.

**Heaviness of a package on its child packages:** For a package denoted as *P*, assume it has \(K\_c\) child packages and the *k*th child is denoted as *A_k_*. Denote \(n\_{1k}\) as the number of strong dependencies of *A_k_*, and \(n\_{2k}\) as the number of strong dependencies of *A_k_* if moving *P* to `Suggests` of *A_k_*, the heaviness of *P* on its child packages denoted as \(h\_c\) is calculated as follows:

\[ h\_c=\frac{1}{K\_c}\sum\_{k=1}^{K\_c}(n\_{1k}-n\_{2k})\]

The heaviness measures the average number of additional dependencies that *P* brings to its child packages.

**Adjusted heaviness on child packages:** Generally, the heaviness on child packages has a trend to decrease with increasing the number of child packages, since it is averaged on the children. To prioritize heavy parents with intermediate numbers of children and to decrease the heaviness values for small \(K\_c\), a penalty term denoted as \(a\) is added to \(K\_c\) as in the following equation where \(h\_c^{adj}\) is the adjusted heaviness for a package on its children. Note \(a\) is set to the same value for all packages and \(a\) is empirically selected to 10.

\[ h\_c^{adj}=\frac{1}{K\_c+a}\sum\_k^{K\_c}(n\_{1k}-n\_{2k})=\frac{K\_c}{K\_c+a}\cdot h\_c \]

Please note, the absolute value of \(h\_c^{adj}\) is meaningless. It is only used for ordering packages.

##### Plots

The following two tabs visualize the distribution of heaviness verse number of child packages. We additionally categorize packages with adjusted heaviness > 30 as highly heavily affecting child packages and with adjusted heaviness between 15 and 30 as medianly heavily affecting child packages.

In the plots, colors (blue to yellow) are mapped to the “density” of the points distribution which are measured as the number of points in a circle with radius of 1% relative to the range both on *x*-axis and *y*-axis.

###### Heaviness on child packages

###### Adjusted heaviness on child packages

##### Table

The highlighted packages in previous plots are listed in the following table. Packages are ordered by adjusted heaviness values. Clicking on the package names will lead to the HTML report of its dependency heaviness analysis. The complete table for all packages is avaiable at https://docs.google.com/spreadsheets/d/1R4ICniRDBuaJlPL6YUN0IzAjmnp73tllPnJeBeLUB98/edit?usp=sharing.

| Package | Repository | Number of child packages | Heaviness on child packages | Adjusted heaviness on child packages |
| --- | --- | --- | --- | --- |
| Rcmdr | CRAN | 47 | 95.9 | 79.1 |
| Seurat | CRAN | 31 | 84.7 | 64.0 |
| RTCGA | Bioconductor | 9 | 135.0 | 63.9 |
| lumi | Bioconductor | 13 | 111.6 | 63.1 |
| brms | CRAN | 8 | 125.8 | 55.9 |
| minfi | Bioconductor | 38 | 57.7 | 45.6 |
| taxize | CRAN | 14 | 77.8 | 45.4 |
| car | CRAN | 183 | 45.9 | 43.6 |
| survminer | CRAN | 24 | 61.6 | 43.5 |
| tidyverse | CRAN | 74 | 49.3 | 43.4 |
| smacof | CRAN | 9 | 89.1 | 42.2 |
| devtools | CRAN | 77 | 46.8 | 41.4 |
| caret | CRAN | 177 | 42.6 | 40.3 |
| GenomicScores | Bioconductor | 25 | 55.6 | 39.7 |
| FactoMineR | CRAN | 51 | 46.4 | 38.8 |
| ggpubr | CRAN | 111 | 42.1 | 38.6 |
| WGCNA | CRAN | 32 | 50.4 | 38.4 |
| drc | CRAN | 18 | 56.5 | 36.3 |
| Deducer | CRAN | 6 | 94.8 | 35.6 |
| AER | CRAN | 21 | 51.7 | 35.0 |
| rms | CRAN | 54 | 39.7 | 33.5 |
| Gviz | Bioconductor | 37 | 41.5 | 32.7 |
| fda | CRAN | 75 | 35.1 | 31.0 |
| Hmisc | CRAN | 250 | 32.0 | 30.8 |
| FlowSOM | Bioconductor | 4 | 107.5 | 30.7 |
| systemfit | CRAN | 15 | 49.3 | 29.6 |
| factoextra | CRAN | 28 | 38.3 | 28.2 |
| DAPAR | Bioconductor | 2 | 168.5 | 28.1 |
| ExperimentHub | Bioconductor | 96 | 30.8 | 27.9 |
| TraMineR | CRAN | 8 | 61.5 | 27.3 |
| qgraph | CRAN | 32 | 35.6 | 27.1 |
| agricolae | CRAN | 12 | 49.5 | 27.0 |
| sampleSelection | CRAN | 5 | 79.4 | 26.5 |
| MESS | CRAN | 6 | 70.5 | 26.4 |
| diffcyt | Bioconductor | 4 | 92.0 | 26.3 |
| DiagrammeR | CRAN | 45 | 31.3 | 25.6 |
| ggbio | Bioconductor | 11 | 48.5 | 25.4 |
| semPlot | CRAN | 7 | 60.3 | 24.8 |
| sparklyr | CRAN | 17 | 38.6 | 24.3 |
| biomod2 | CRAN | 2 | 145.5 | 24.2 |
| doBy | CRAN | 14 | 41.5 | 24.2 |
| rstanarm | CRAN | 6 | 63.8 | 23.9 |
| flowDensity | Bioconductor | 3 | 101.0 | 23.3 |
| pec | CRAN | 7 | 55.9 | 23.0 |
| fields | CRAN | 174 | 24.1 | 22.8 |
| randomForestSRC | CRAN | 7 | 54.3 | 22.4 |
| flexsurv | CRAN | 16 | 36.2 | 22.3 |
| klaR | CRAN | 11 | 42.5 | 22.3 |
| beadarray | Bioconductor | 7 | 54.0 | 22.2 |
| xcms | Bioconductor | 16 | 35.9 | 22.1 |
| GSVA | Bioconductor | 18 | 34.3 | 22.0 |
| missMDA | CRAN | 8 | 48.8 | 21.7 |
| biodb | Bioconductor | 5 | 64.2 | 21.4 |
| MSnbase | Bioconductor | 27 | 29.1 | 21.3 |
| sva | Bioconductor | 38 | 26.9 | 21.3 |
| clusterProfiler | Bioconductor | 36 | 27.1 | 21.2 |
| golem | CRAN | 19 | 31.6 | 20.7 |
| rstan | CRAN | 119 | 22.5 | 20.7 |
| ensembldb | Bioconductor | 27 | 28.2 | 20.6 |
| ez | CRAN | 10 | 40.7 | 20.4 |
| VariantAnnotation | Bioconductor | 71 | 22.8 | 20.0 |
| matlib | CRAN | 4 | 69.5 | 19.9 |
| ecospat | CRAN | 1 | 216.0 | 19.6 |
| GenomicFeatures | Bioconductor | 186 | 20.3 | 19.3 |
| forecast | CRAN | 116 | 20.9 | 19.3 |
| DESeq2 | Bioconductor | 93 | 20.8 | 18.8 |
| mapview | CRAN | 11 | 35.8 | 18.8 |
| MLInterfaces | Bioconductor | 5 | 55.8 | 18.6 |
| BSgenome | Bioconductor | 230 | 19.3 | 18.5 |
| testthat | CRAN | 120 | 19.8 | 18.3 |
| haplo.stats | CRAN | 5 | 54.6 | 18.2 |
| photobiology | CRAN | 10 | 36.2 | 18.1 |
| MplusAutomation | CRAN | 12 | 32.6 | 17.8 |
| RTN | Bioconductor | 3 | 75.7 | 17.5 |
| kableExtra | CRAN | 65 | 20.1 | 17.4 |
| phyloseq | Bioconductor | 23 | 24.7 | 17.2 |
| AnnotationDbi | Bioconductor | 722 | 17.4 | 17.1 |
| TFBSTools | Bioconductor | 8 | 38.4 | 17.1 |
| mosaic | CRAN | 7 | 40.7 | 16.8 |
| biovizBase | Bioconductor | 9 | 35.2 | 16.7 |
| methylPipe | Bioconductor | 2 | 99.0 | 16.5 |
| biomaRt | Bioconductor | 97 | 18.0 | 16.4 |
| HH | CRAN | 6 | 43.7 | 16.4 |
| spdep | CRAN | 59 | 19.2 | 16.4 |
| SpatialExperiment | Bioconductor | 9 | 34.4 | 16.3 |
| fdapace | CRAN | 9 | 34.3 | 16.3 |
| ChemmineR | Bioconductor | 11 | 30.9 | 16.2 |
| MDSMap | CRAN | 2 | 97.5 | 16.2 |
| Ecdat | CRAN | 4 | 56.2 | 16.1 |
| affycoretools | Bioconductor | 1 | 176.0 | 16.0 |
| ibd | CRAN | 2 | 96.0 | 16.0 |
| dvmisc | CRAN | 4 | 56.0 | 16.0 |
| NMF | CRAN | 21 | 23.4 | 15.9 |
| mixOmics | Bioconductor | 14 | 27.0 | 15.8 |
| ENMeval | CRAN | 3 | 68.0 | 15.7 |
| fda.usc | CRAN | 12 | 28.3 | 15.5 |
| spm | CRAN | 1 | 171.0 | 15.5 |
| geojsonio | CRAN | 9 | 32.3 | 15.3 |
| CCA | CRAN | 4 | 53.2 | 15.2 |
| JWileymisc | CRAN | 2 | 91.0 | 15.2 |
| leaflet | CRAN | 77 | 17.1 | 15.2 |
| GEOquery | Bioconductor | 26 | 20.8 | 15.1 |
| plsRglm | CRAN | 3 | 65.3 | 15.1 |
| MethylAid | Bioconductor | 1 | 165.0 | 15.0 |
| hdrcde | CRAN | 11 | 28.7 | 15.0 |
