## Supplementary File 3 for "*pkgndep*: a tool for analyzing dependency heaviness of R packages": suppl_3_heaviness_on_children.Rmd

````
---
title: "Heaviness on child packages"
author: "Zuguang Gu"
date: '`r Sys.Date()`'
output:
  html_document:
    fig_caption: true
---

```{r, echo = FALSE}
library(knitr)
knitr::opts_chunk$set(
    warning = FALSE,
    echo = FALSE,
    fig.align = "center")
```

### Definition

**Strong parent packages:** The packages listed in the <code>Depends</code>, <code>Imports</code>
and <code>LinkingTo</code> fields of the <code>DESCRIPTION</code> of a package denoted as _P_.
They are also called the strong direct dependency packages of _P_. Strong parent
packages are enforced to be installed when installing package _P_. To make it
easy to discuss, in the text we always refer parent packages to strong parent
packages.

**Strong dependency packages:** The total packages by recursively looking for
parent packages. They are also called upstream packages. Note strong dependency packages
contain parent packages. Strong dependency packages are enforced to be
installed when installing package _P_.

**Heaviness of a package on its child packages:** For a package denoted as _P_, assume
it has $K_c$ child packages and the <i>k</i>th child is denoted as _A<sub>k</sub>_. Denote $n_{1k}$ as the
number of strong dependencies of _A<sub>k</sub>_, and $n_{2k}$ as the number of strong
dependencies of _A<sub>k</sub>_ if moving _P_ to <code>Suggests</code> of _A<sub>k</sub>_, the heaviness of _P_ on its
child packages denoted as $h_c$ is calculated as follows:

$$ h_c=\frac{1}{K_c}\sum_{k=1}^{K_c}(n_{1k}-n_{2k})$$

The heaviness measures the average number of additional dependencies that _P_
brings to its child packages.

**Adjusted heaviness on child packages:** Generally, the heaviness on child
packages has a trend to decrease with increasing the number of child packages,
since it is averaged on the children. To prioritize heavy parents with
intermediate numbers of children and to decrease the heaviness values for
small $K_c$, a penalty term denoted as $a$ is added to $K_c$ as in the
following equation where $h_c^{adj}$ is the adjusted heaviness for a package
on its children. Note $a$ is set to the same value for all packages and $a$ is
empirically selected to 10.

$$ h_c^{adj}=\frac{1}{K_c+a}\sum_k^{K_c}(n_{1k}-n_{2k})=\frac{K_c}{K_c+a}\cdot h_c $$

Please note, the absolute value of $h_c^{adj}$ is meaningless. It is only used for ordering packages.

### Plots {.tabset}

The following two tabs visualize the distribution of heaviness verse number of child packages. We additionally
categorize packages with adjusted heaviness > 30 as highly heavily affecting child packages and with adjusted heaviness between 15 and 30
as medianly heavily affecting child packages.

In the plots, colors (blue to yellow) are mapped to the "density" of the points distribution which are measured as the number
of points in a circle with radius of 1% relative to the range both on _x_-axis and _y_-axis.

#### Heaviness on child packages

```{r, fig.width = 10, fig.height = 5, echo = FALSE}
library(pkgndep)
library(ggplot2)
library(ggnewscale)

df = load_pkg_stat_snapshot()
df = df[df$n_children > 0,]

CUTOFF = list()
CUTOFF$adjusted_heaviness_on_children = c(15, 30)

heaviness_cate = ifelse(df$adjusted_heaviness_on_children >= CUTOFF$adjusted_heaviness_on_children[2], "high", ifelse(df$adjusted_heaviness_on_children >= CUTOFF$adjusted_heaviness_on_children[1], "median", "low"))
df$heaviness_cate = heaviness_cate
heaviness_color = c("high" = "red", "median" = "orange")
repo = ifelse(grepl("bioconductor", df$repository), "Bioconductor", "CRAN")
df$repo = factor(repo, levels = c("CRAN", "Bioconductor"))

x = log10(df$n_children)
y = df$heaviness_on_children
max_x = max(x)
max_y = max(y)
x = x/max(x)
y = y/max(y)

df$n_neighbours = 0
for(r in c("CRAN", "Bioconductor")) {
	l = df$repo == r
	d = dist(cbind(x[l], y[l]))
	d = as.matrix(d)
	df$n_neighbours[l] = apply(d, 1, function(x) sum(x < 0.001))
}
df = df[order(df$n_neighbours), ] # put points with higher n_neighbours on top

l_median = df$adjusted_heaviness_on_children >= CUTOFF$adjusted_heaviness_on_children[1] & df$adjusted_heaviness_on_children < CUTOFF$adjusted_heaviness_on_children[2]
l_high = df$adjusted_heaviness_on_children >= CUTOFF$adjusted_heaviness_on_children[2]
l_low = !(l_median | l_high)
ggplot(df[l_low, ], aes(n_children, heaviness_on_children, color = log10(n_neighbours))) +
    geom_point(size = 0.5) + scale_color_viridis_c(breaks = c(0, 1, 2, 3), labels = c(1, 10, 100, 1000), name = "Number of\nneighbor points") +
    scale_x_continuous(trans='log10') +
    new_scale_color() +
    geom_point(aes(n_children, heaviness_on_children, color = "median"), data = df[l_median, ]) +
    geom_point(aes(n_children, heaviness_on_children, color = "high"), data = df[l_high, ]) +
    scale_color_manual(name = "Heaviness category", values = heaviness_color) +
    ggrepel::geom_text_repel(aes(n_children, heaviness_on_children, label = package), data = df[l_high, ], min.segment.length = 0, box.padding = 0.5, max.overlaps = Inf, show.legend = FALSE, size = 3, color = "red") +
    labs(x = "Number of child packages", y = "Heaviness on child packages") +
    ggtitle("Heaviness on child packages") +
    facet_wrap(vars(repo))
```

#### Adjusted heaviness on child packages


```{r, fig.width = 10, fig.height = 5, echo = FALSE}
x = log10(df$n_children)
y = df$adjusted_heaviness_on_children
max_x = max(x)
max_y = max(y)
x = x/max(x)
y = y/max(y)

df$n_neighbours = 0
for(r in c("CRAN", "Bioconductor")) {
	l = df$repo == r
	d = dist(cbind(x[l], y[l]))
	d = as.matrix(d)
	df$n_neighbours[l] = apply(d, 1, function(x) sum(x < 0.001))
}
df = df[order(df$n_neighbours), ] # put points with higher n_neighbours on top

l_median = df$adjusted_heaviness_on_children >= CUTOFF$adjusted_heaviness_on_children[1] & df$adjusted_heaviness_on_children < CUTOFF$adjusted_heaviness_on_children[2]
l_high = df$adjusted_heaviness_on_children >= CUTOFF$adjusted_heaviness_on_children[2]
l_low = !(l_median | l_high)
ggplot(df[l_low, ], aes(n_children, adjusted_heaviness_on_children, color = log10(n_neighbours))) +
    geom_point(size = 0.5) + scale_color_viridis_c(breaks = c(0, 1, 2, 3), labels = c(1, 10, 100, 1000), name = "Number of\nneighbor points") +
    scale_x_continuous(trans='log10') +
    new_scale_color() +
    geom_point(aes(n_children, adjusted_heaviness_on_children, color = "median"), data = df[l_median, ]) +
    geom_point(aes(n_children, adjusted_heaviness_on_children, color = "high"), data = df[l_high, ]) +
    scale_color_manual(name = "Heaviness category", values = heaviness_color) +
    ggrepel::geom_text_repel(aes(n_children, adjusted_heaviness_on_children, label = package), data = df[l_high, ], min.segment.length = 0, box.padding = 0.5, max.overlaps = Inf, show.legend = FALSE, size = 3, color = "red") +
    labs(x = "Number of child packages", y = "Adjusted heaviness on child packages") +
    ggtitle("Adjusted heaviness on child packages") +
    facet_wrap(vars(repo))
```

### Table

The highlighted packages in previous plots are listed in the following table. Packages are ordered by adjusted heaviness values. Clicking on
the package names will lead to the HTML report of its dependency heaviness analysis. The complete table for all packages is avaiable at https://docs.google.com/spreadsheets/d/1R4ICniRDBuaJlPL6YUN0IzAjmnp73tllPnJeBeLUB98/edit?usp=sharing.

<style>
.heaviness-high {
	padding: 2px 4px;
	background-color: red;
	border-radius:  4px;
}

.heaviness-high a {
	color: white;
}

.heaviness-median {
	padding: 2px 4px;
	background-color: orange;
	border-radius:  4px;
}

.heaviness-median a {
	color: white;
}
</style>

```{r}
library(kableExtra)
library(GetoptLong)
tb = df[l_median | l_high, ]
tb$heaviness_on_children = round(tb$heaviness_on_children, 1)
tb$adjusted_heaviness_on_children = round(tb$adjusted_heaviness_on_children, 1)
tb = tb[order(tb$adjusted_heaviness_on_children, decreasing = TRUE), ]

tb$package = qq("<span class='heaviness-@{tb$heaviness_cate}'><a href='https://pkgndep.github.io/prefix_@{tolower(substr(tb$package, 1, 1))}/@{tb$package}_dependency_report.html'>@{tb$package}</a></span>", collapse = FALSE)

tb = tb[, c("package", "repo", "n_children", "heaviness_on_children", "adjusted_heaviness_on_children")]
colnames(tb) = c("Package", "Repository", "Number of child packages", "Heaviness on child packages", "Adjusted heaviness on child packages")
kbl(tb, escape = FALSE, row.names = FALSE) %>%
  kable_styling()
```
````
