## Supplementary File 4 for "*pkgndep*: a tool for analyzing dependency heaviness of R packages": suppl_4_cola.html

Dependency heaviness analysis for package cola


### Dependency heaviness analysis for package cola

###### Zuguang Gu

#### 2022-02-02

*cola* is a package for consensus partitioning analysis. It implements comprehensive functionalities not only for consensus partitioning, but also for downstream anlaysis such as dimension reduction analysis, signature analysis and functional enrichment analysis. It integrates a lot of other packages. The dependency heaviness analysis is as follows:

```
library(pkgndep)
x = pkgndep("cola")
```

```
plot(x)
```

You can drag the plot into a new tab if it is too small to read. The dependency analysis for *cola* is also available at https://pkgndep.github.io/prefix\_c/cola\_dependency\_report.html.

The dependency heaviness analysis shows the number of total dependency for *cola* is 248, which means, if the full functionality of *cola* is required by a user, he or she needs to install all 248 upstream packages. *cola* performs consensus partitioning as its core analysis which is expected to be very frequently used by users, while other downstream analysis such as functional enrichment analysis are less used. On the other hand, dependency packages for downstream analysis contribute very high heaviness to *cola*. For example, package *clusterPrifiler* which is for functional enrichment analysis contribute a heaviness of 91 and package *ReactomePA* which provides Reactome pathways for enrichment analysis contribute a heaviness of 96. Since we are also the developers of package *cola*, we arranged the parents of *cola* in a way that only package related to the core analysis were put as strong parents, while those for secondary analysis were put as weak parents. This makes the number of strong dependencies of *cola* reduced to only 64.
