## Supplementary File 5 for "*pkgndep*: a tool for analyzing dependency heaviness of R packages": suppl_5_co_heaviness.html

Co-heaviness of two parent packages


### Co-heaviness of two parent packages

###### Zuguang Gu

#### 2022-02-02

The heaviness measures the number of additional dependency packages that a parent package imports, which are the dependency packages not imported by any of the other parent packages. However, there are scenarios when multiple parents import similar sets of dependencies, which results in heaviness for individual parent very small.

We take the **DESeq2** package as an example. First we generate the dependency heatmap.

```
library(pkgndep)
x = pkgndep("DESeq2")
```

```
plot(x)
```

You can drag the plot into a new tab if it is too small to read. The dependency analysis for *DESeq2* is also available at https://pkgndep.github.io/prefix\_d/DESeq2\_dependency\_report.html.

As can be observed from the heatmap, **DESeq2**’s two parents **geneplotter** and **genefilter** (the last two rows in “Import” category in the heatmap) import 51 and 53 dependency packages, among them 50 packages are the same. Due to the high overlap, the heaviness of **geneplotter** and **genefilter** on **DESeq2** are only 1 and 2 respectively. However, if taking the two parents together, i.e., by moving both parents to “Suggests” of **DESeq2**, 23 dependency packages can be reduced.

Here we define **the co-heaviness** that measures the number of additional dependency packages brought by two parent packages. Denote a package as *P* and its two strong parent packages as *A* and *B*, i.e., parent packages in “Depends”, “Imports” and “LinkingTo”, denote \(S\_A\) as the set of reduced dependency packages when only moving *A* to “Suggests” of *P*, denote \(S\_B\) as the set of reduced dependency packages when only moving *B* to “Suggests” of *P*, and denote \(S\_{AB}\) as the set of reduced dependency packages when moving *A* and *B* together to “Suggests” of *P*, the co-heaviness of *A*, *B* on *P* is calculatd as

\[ \left | S\_{AB} \setminus \cup (S\_A, S\_B) \right | \]

which is the number of reduced package only caused by co-action of A and B. Symbol \(A \setminus B\) is the set of elements in *A* not in *B* and \(|A|\) is the number of elements in set *A*.

**pkgndep** provides a function `co_heaviness()` that calculates co-heaviness of two parent packages. It returns a co-heaviness matrix. Please note it only returns the matrix for strong parents.

```
m = co_heaviness(x)
m
```

```
##                      S4Vectors IRanges GenomicRanges SummarizedExperiment
## S4Vectors                    0       0             0                    0
## IRanges                      0       0             0                    0
## GenomicRanges                0       0             0                    1
## SummarizedExperiment         0       0             1                    4
## methods                      0       0             0                    0
## stats4                       0       0             0                    0
## Rcpp                         0       0             0                    0
## BiocGenerics                 0       0             0                    0
## Biobase                      0       0             0                    0
## locfit                       0       0             0                    0
## BiocParallel                 0       0             0                    0
## ggplot2                      0       0             0                    0
## geneplotter                  0       0             0                    0
## genefilter                   0       0             0                    0
## RcppArmadillo                0       0             0                    0
##                      methods stats4 Rcpp BiocGenerics Biobase locfit
## S4Vectors                  0      0    0            0       0      0
## IRanges                    0      0    0            0       0      0
## GenomicRanges              0      0    0            0       0      0
## SummarizedExperiment       0      0    0            0       0      0
## methods                    0      0    0            0       0      0
## stats4                     0      0    0            0       0      0
## Rcpp                       0      0    0            0       0      0
## BiocGenerics               0      0    0            0       0      0
## Biobase                    0      0    0            0       0      0
## locfit                     0      0    0            0       0      1
## BiocParallel               0      0    0            0       0      0
## ggplot2                    0      0    0            0       0      0
## geneplotter                0      0    0            0       0      0
## genefilter                 0      0    0            0       0      0
## RcppArmadillo              0      0    0            0       0      0
##                      BiocParallel ggplot2 geneplotter genefilter RcppArmadillo
## S4Vectors                       0       0           0          0             0
## IRanges                         0       0           0          0             0
## GenomicRanges                   0       0           0          0             0
## SummarizedExperiment            0       0           0          0             0
## methods                         0       0           0          0             0
## stats4                          0       0           0          0             0
## Rcpp                            0       0           0          0             0
## BiocGenerics                    0       0           0          0             0
## Biobase                         0       0           0          0             0
## locfit                          0       0           0          0             0
## BiocParallel                    8       0           0          0             0
## ggplot2                         0      21           1          1             0
## geneplotter                     0       1           1         23             0
## genefilter                      0       1          23          2             0
## RcppArmadillo                   0       0           0          0             1
```

The co-heaviness matrix can be visualized as a heatmap.

```
library(ComplexHeatmap)
Heatmap(m, name = "co-heaviness")
```

There are two major reasons for the high co-heaviness from two parent packages. 1. Parent A also depends on package B, thus all the dependency packages brought by package B will also be included in the dependency packages from package A. E.g., package **ffpe**’s two parents **lumi** and **methylumi** where **lumi** also depends on **methylumi** and most of the heaviness comes from **methylummi**. 2. Although A and B do not depend on each other, they have a common upstream package, which brings the same dependencies to A and B. Such as the example of **DESeq2**, its two parent packages **geneplotter** and **genefilter** have a common upstream pacakge **annotate** that contribute many dependencies to both package.

Finally, co-heaviness captures additional dependency packages from and only from two parents, while it does work for more parents. However, the effect of heaviness from multiple parents can always be easily observed from the dependency heatmap.
